# Multi-omics analysis reveals novel interplays between intratumoral bacteria and glioma

**DOI:** 10.1101/2023.10.08.561332

**Authors:** Ting Li, Zhanyi Zhao, Meichang Peng, Cheng Wang, Feiyang Luo, Meiqin Zeng, Kaijian Sun, Zhencheng Fang, Yunhao Luo, Qiyuan Huang, Yugu Xie, Jiaxuan Wang, Jian-Dong Huang, Hongwei Zhou, Haitao Sun

## Abstract

Emerging evidence highlights the potential impact of intratumoral microbiota on cancer. However, the microbial composition and function in glioma remains elusive. Consequently, our study aimed to investigate the microbial community composition in glioma tissues and elucidate its role in glioma development. We parallelly performed microbial profiling, transcriptome sequencing and metabolomics detection on tumor and adjacent normal brain tissues obtained from 50 glioma patients. We employed immunohistochemistry, multicolor immunofluorescence and FISH staining to observe the presence and location of bacteria. Furthermore, an animal model was employed to validate the impact of key bacteria on glioma development. Six genera were found to be significantly enriched in glioma tissues compared to its adjacent normal brain tissues, including *Fusobacterium*, *Longibaculum*, *Intestinimonas*, *Pasteurella*, *Limosilactobacillus* and *Arthrobacter*. Both bacterial RNA and LPS were observed in glioma tissues. Multicolor immunofluorescence analysis showed higher bacterial LPS levels in tumor cells than in macrophages and in glioma tissue than in adjacent normal brain tissue. Integrated microbiomics, transcriptomics, and metabolomics revealed that genes associated with intratumoral microbes were enriched in multiple synapse-associated pathways and that metabolites associated with intratumoral microbes were (R)-N-methylsalsolinol, N-acetylaspartylglutamic acid, and N-acetyl-L-aspartic acid. Further mediation analysis suggested that intratumoral microbiome may affect the expression of neuron-related genes through bacteria-associated metabolites. In addition, a glioma mouse model suggested that *Fusobacterium nucleatum* promoted glioma growth by increasing the levels of N-acetylneuraminic acid and the expression levels of CCL2, CXCL1, and CXCL2. In conclusion, our findings shed light on the intricate interplays between intratumoral bacteria and glioma.

## Introduction

Glioma, as the predominant primary malignancy of the central nervous system[1], poses a formidable challenge for treatment worldwide owing to its high recurrence, mortality, and poor prognosis. It is imperative to further elucidate the pathogenic mechanisms of glioma and develop novel therapeutic strategies. Increasing evidence has underscored the significant contributions made by intratumoral microbiota in tumor progression[2], metastasis[3] and treatment[4], with a high likelihood of serving as biomarkers for early diagnosis and treatment of tumors.

Whether microbiota exists in glioma, or even in the brain, remains issues under debate. The brain has traditionally been regarded as a sterile organ due to the blood-brain barrier. However, recent evidence indicates that microbiota may directly inhabit the brain under non-inflammatory and non-traumatic conditions. Gram-negative bacterial molecules and *Porphyromonas gingivalis* were detected in the brains of Alzheimer’s disease patients and found to be associated with tau protein accumulation.[5–7]. Rombert et al. [8] reported in an abstract at the American Neuroscience Annual Meeting that they found rod-shaped bacteria in the healthy human post-mortem brains under electron microscopy.

In addition, a study conducted a comprehensive detection and analysis of the intratumoral microbiota of seven types of solid tumors, including glioblastoma[9]. They found that glioblastoma harbored a unique microbial community and that the level of bacterial DNA was not low. Due to the small sample size and the limitations of the experimental design, this study did not reveal the role of intratumoral microbiota in glioblastoma.

In our previous work[10], we used intact tissues and combined tissue clearing, immunofluorescence staining methods to observe bacterial lipopolysaccharides (LPS) in glioma in three-dimensional space. Although we further confirmed the presence of bacterial LPS in human glioma tissue morphologically, more evidence is needed to elucidate the spatial relationship and function of bacteria in glioma.

The study of the tumor microbiome is a challenging task due to the low intratumor bacterial biomass and uncultivability of some microbial species. Novel high-resolution techniques are urgently required to facilitate further research. Multi-omics techniques can elucidate tumor microbiota at different levels, thereby facilitating a comprehensive understanding of the intricate biological processes associated with tumor microbiota.

To this end, we simultaneously performed 16S rRNA sequencing, metabolomics and transcriptomics analyses on glioma tissue (G) samples and adjacent normal brain tissue (NAT) samples. Our results suggested that intratumoral microbiota of glioma may affect the expression of neuron-related genes through bacteria-associated metabolites.

## Materials and Method

### Human subjects and sample collection

We retrospectively obtained 110 fresh frozen tissues, 8 paraffin tissues and 10 stool samples from glioma patients using strict eligibility criteria. The clinical information of the patients was shown in Additional file 1. The stool samples were taken before cancer treatment, and individuals who received preoperative radiation or chemotherapy treatment or had a previous history of glioma were excluded. Allsamples were collected from Zhujiang Hospital of Southern Medical University (Guangzhou, China).

Tissue samples were meticulously collected and promptly processed in the operating room to minimize the risk of contamination. Samples were collected in sterile cryotubes and transported using a portable liquid nitrogen tank to the Clinical Biobank Center, where they were kept at -196°C for long-term storage until DNA extraction. Fecal samples are rapidly transferred to a -80 °C freezer for storage until further use.

### Mice

Athymic BALB/c nude mice (4 weeks; 20–25 g) were purchased from GemPharmatech Corporation (Nanjing, China) and maintained in a specific pathogen-free environment under a 12-h light-dark cycle with free access to food and water.

### Multiplex Immunofluorescent Assay

4 μM paraffin sections were deparaffinized in xylene and rehydrated in a series of graded alcohols. Antigen extraction was performed in citrate buffer (pH 6), boiled at high power in a microwave oven for 20 seconds, maintained at low power for 5 minutes at low boiling state, and cooled naturally after turning off the heat. Multiplex fluorescence labeling was performed using TSA-dendron-fluorophores with NEON 7-color Allround Discovery Kit for FFPE (Histova Biotechnology). Multiplex antibody panels applied in this study are: CD68 (abcam#213363, 1:200), GFAP (abcam#68428, 1:200), LPS(Lipopolysaccharide Core, HycultBiotech#HM6011,1:200), DAPI (abcam#ab104139). After detecting all antibodies, images were taken using TissueFAXS imaging software (v7.134), and viewed with TissueFAXS Viewer software, and fluorescent positive cells were counted using StrataQuest tissue flow cytometry quantitative analysis system. For detailed analytical methods, please see Additional file 2.

### Immunohistochemistry

The pretreatment of paraffin tissue sections was performed according to the steps of multicolor immunofluorescence. The primary antibodies used are LPS (HycultBiotech#HM6011,1:1000) and LTA (GeneTex#16470,1:1000). The dyes used are DAB kits (Servicebio#G1211). Slides were scanned with Pannoramic Scan II (3D HISTECH) and images were generated with SlideViewer (v2.5, 3D HISTECH).

### 16S rRNA FISH

The 4 μm FFPE tissue slides were routinely deparaffinized and hydrated. Slides were stained for bacterial 16S rRNA (Cy3-labeled EUB338 probes, Future Biotech #FBFPC001) or negative control (Cy3-labeled nonspecific complement probe, Future Biotech #FBFPC001) using the direct fluorescent bacteria in situ hybridization detection kit (Future Biotech #FB0016) according to the manufacturer’s instructions. Slides were scanned with Pannoramic Scan II (3D HISTECH) and images were generated with SlideViewer (v2.5, 3D HISTECH).

### DNA extraction and sequencing

Microbial DNA was extracted from tissue samples using the QIAGEN DNeasy PowerSoil Kit (#47014) according to manufacturer’s protocol. Library construction and sequencing were performed by Novogene Corporation (Beijing, China). The downstream data processing was performed using EasyAmplicon (v1.12), VSEARCH (v2.15.2) and USEARCH (v10.0.240). For detailed analytical methods, please see Additional file 2.

### Microbiome data analysis

The vegan package (v2.5-6) in R (v4.1) was performed for alpha diversity analysis. The unweighted UniFrac distance matrix was generated by USEARCH. Beta diversity was calculated using principal coordinate analysis (PCoA). To visualize the results of diversity analysis, the R software package ggplot2 was used. The compositions of the microbial community in two groups were presented as stacked bar plots at the phylum levels. Analysis of variance was performed with the software package STAMP (v 2.1.3) using the storey false-discovery rate approach for correction of P values. LEfSe was performed using an online utility (http://www.ehbio.com/ImageGP/index.php/Home/Index/LEFSe.html) to analyze the differences of bacterial abundance at different bacterial classification levels. Finally, Phylogenetic Investigation of Communities by Reconstruction of Unobserved States (PICRUSt) was used to predict microbial functional signatures. In the PICRUSt analysis, the significant KEGG pathways (level 3) among species were analyzed via Welch’s t-test in STAMP using the Bonferroni correction. Statistical analysis was conducted with Graghpad Prism (v8.3) software.

### Contamination correction

To prevent a false-positive rate, 15 negative controls, including 5 environmental controls, 5 DNA extraction controls, and 5 PCR controls, were prepared and sequenced alongside the tissue samples. In order to correct for contamination, binomial tests were conducted between samples and negative controls to determine their abundance. In negative control samples, the frequency of occurrence of a taxon was used to estimate p of a binomial distribution. For the binomial test, x and n represent the number of occurrences and totals, respectively. In the following analysis, those taxa with a *P*-value of 0.05 were kept.

### mRNA-seq

Frozen human glioma tissues, adjacent normal brain tissue and mice tumor tissues were used for RNA extraction. A profiler service provided by Genergy Biotechnology Corporation (Shanghai, China) was used for RNA-seq. DESeq2 Bioconductor package was used to identify differentially expressed genes (DEGs), and remarkable DEGs were selected based on *P* < 0.05, *FDR* < 0.05, and a log2 (fold change) > 1. The DEGs were also submitted for Kyoto Encyclopedia of Genes and Genomes (KEGG) analysis, and significant pathways with *P* < 0.05 were shown. For detailed analytical methods, please see Additional file 2.

### Metabolomics

Frozen human glioma tissues, adjacent normal brain tissue and mice tumor tissues were used for metabolomic profiling. Untargeted metabolomics profiling was performed by Biotree Biotechnology Corporation (Shanghai, China). All data were analyzed by LC-MS/MS on UHPLC system. The differential metabolites were screened by combining the results of the student’s t-test (*P* < 0.05) and the Variable Importance in the Projection (VIP > 1) of the first principal component of the OPLS-DA model. Using the Kyoto Encyclopedia of Genes and Genomes Pathway database, we performed KEGG annotation of differential metabolites. By mapping the differential metabolites to authoritative databases such as KEGG, PubChem, HMDB, after obtaining the matching information of the differential metabolites, we searched the human pathway databases and analyzed the metabolic pathways. For detailed analytical methods, please see Additional file 2.

### Multi-omics analyses

Correlation analysis of differentially expressed genes, differential metabolites, and differential bacteria was performed calculating the Spearman correlation coefficient using package psych in R. Subsequently, we constructed a network integrating the interactions among differentially expressed genes, metabolites, and bacteria, and visualized it using Cytoscape software. Mediation analysis was carried out using the mediation package.

### Fusobacterium nucleatum culture

Fusobacterium nucleatum ATCC 25586 (Fn) was gifted kindly by Dr. Songhe Guo. Fn was grown anaerobically at 37 °C for 48-72 h in blood agar plate (Oxoid, UK). Fn were centrifuged and then suspended to a concentration of 1×10^6^ colony-forming units (CFUs)/ml with PBS for the animal experiment.

### Animal experiments

U87 cells (1×10^6^) in the logarithmic growth phase in 100 μL of PBS were subcutaneously injected into the right flank of the mice. When the tumor volume reached 200 mm^3^ (day 1), mice were randomly divided into four treatment groups (intratumoral injection): PBS group received injection of PBS, Fn group received injection of the 5×10^6^ CFU *Fusobacterium nucleatum* (Fn), Fn+MTZ group received injection of Fn and metronidazole (MTZ) treatment in addition, and MTZ group received injection of MTZ alone. For the Fn group and Fn+MTZ group, the bacteria were initially injected with PBS or metronidazole for 30 minutes before inoculation. On days 2 and 3, treat with metronidazole or a PBS vehicle twice daily, at least 8 hours apart. on days 4-7, once a day. On day 8, tumor tissue was obtained and weighed. Tumor tissue was immediately frozen in liquid nitrogen for subsequent experiments.

## Results

### Study design

To investigate tumor-associated microbial community, we enrolled 50 glioma patients in this study. Fifty glioma and 15 matched adjacent normal brain fresh frozen tissues were used for 16S rRNA gene sequencing. We simultaneously collected 15 negative control samples, including 5 environmental controls, 5 DNA extraction controls, and 5 PCR controls for contamination filtering. Twenty glioma and six matched adjacent normal brain fresh frozen tissues were used for untargeted metabolomics analysis. Sixteen glioma and three matched adjacent normal brain fresh frozen tissues were used for transcriptomics sequencing. Four glioma and four adjacent normal brain paraffin tissues were used for bacterial imaging. To further elucidate the role of *Fusobacterium nucleatum*, the most renowned tumor-associated species within the *Fusobacterium* genus, we designed an in vivo animal model experiment. Schematic diagram of the design of the entire study was shown in Fig. 1.

**Fig. 1.**
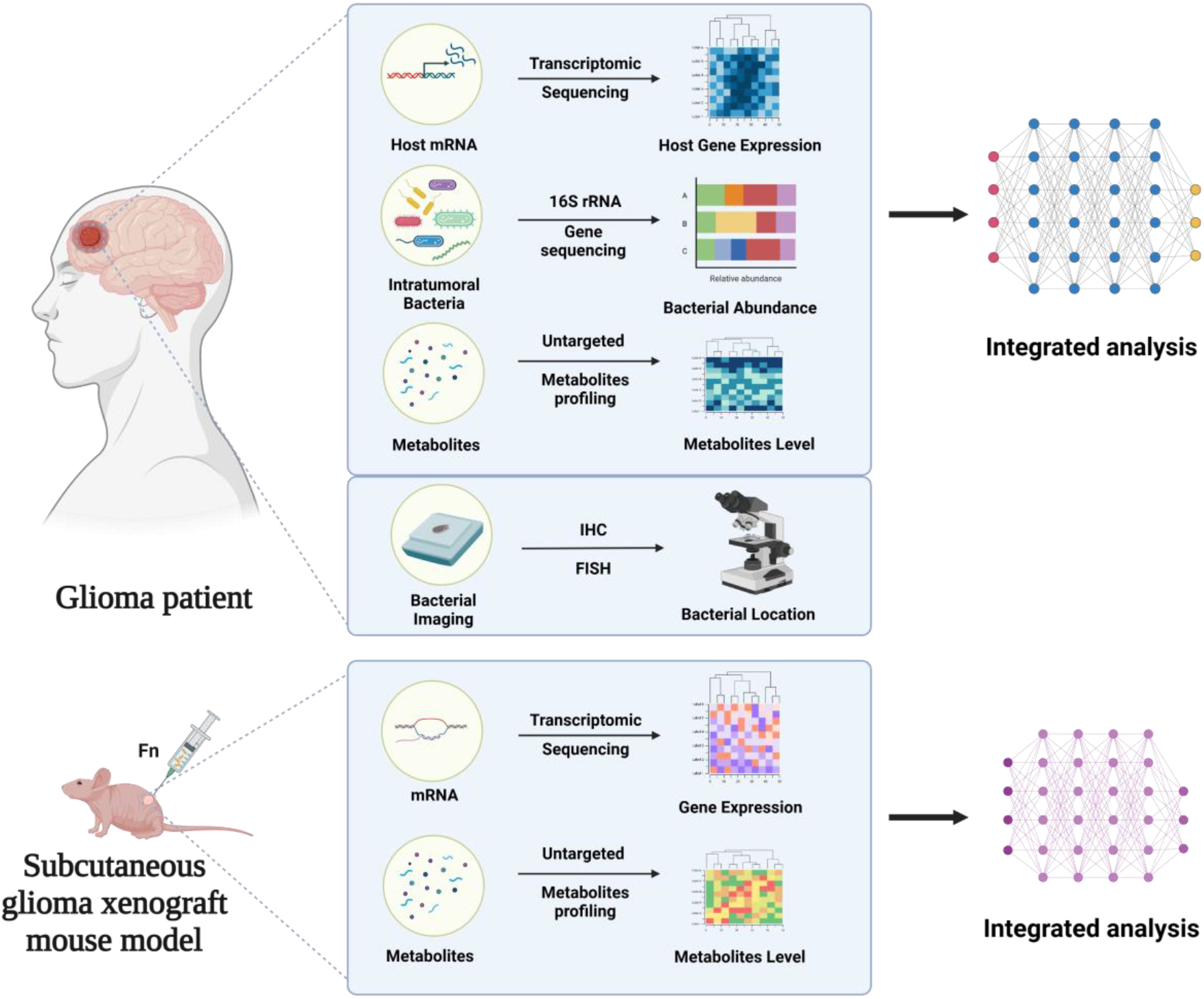
Schematic of research design.

### Profiling of microbiota in human glioma tissues

To determine the homogeneity of the microbiota within glioma tissues, we conducted a survey of microbial diversity in 35 unmatched tissues and 15 matched tissues, and found no significant difference in α and β diversity between the two groups. Therefore, in the subsequent analysis, we selected 50 glioma samples as representatives of the glioma group (Additional file 3: Fig. S1a, b)

In our present microbiome investigation, overall alpha diversity of the G group was significantly higher than that of the NAT group (Fig. 2a). Employing the unconstrained PCoA with Bray-Curtis distance analysis, we demonstrated a clear separation between the glioma-associated microbiota and those present in adjacent normal brain tissue (Fig. 2b). The results showed the microbial composition between G and NAT was markedly different. Specifically, we found that at the phylum level, the tumor-associated microbiota was dominated by *Firmicutes* and *Proteobacteria*, followed by *Actinobacteria*, *Fusobacteria* and *Bacteroidetes* (Fig. 2c). The relative abundance of the phyla *Firmicutes* and *Fusobacteria* was greater in the G group than in the NAT group, whereas the *Proteobacteria* phylum exhibited an inverse relationship (Fig. 2d). These observations align with previous reports on investigations of the potential brain microbiome in Alzheimer’s disease.[11]

**Fig. 2.**
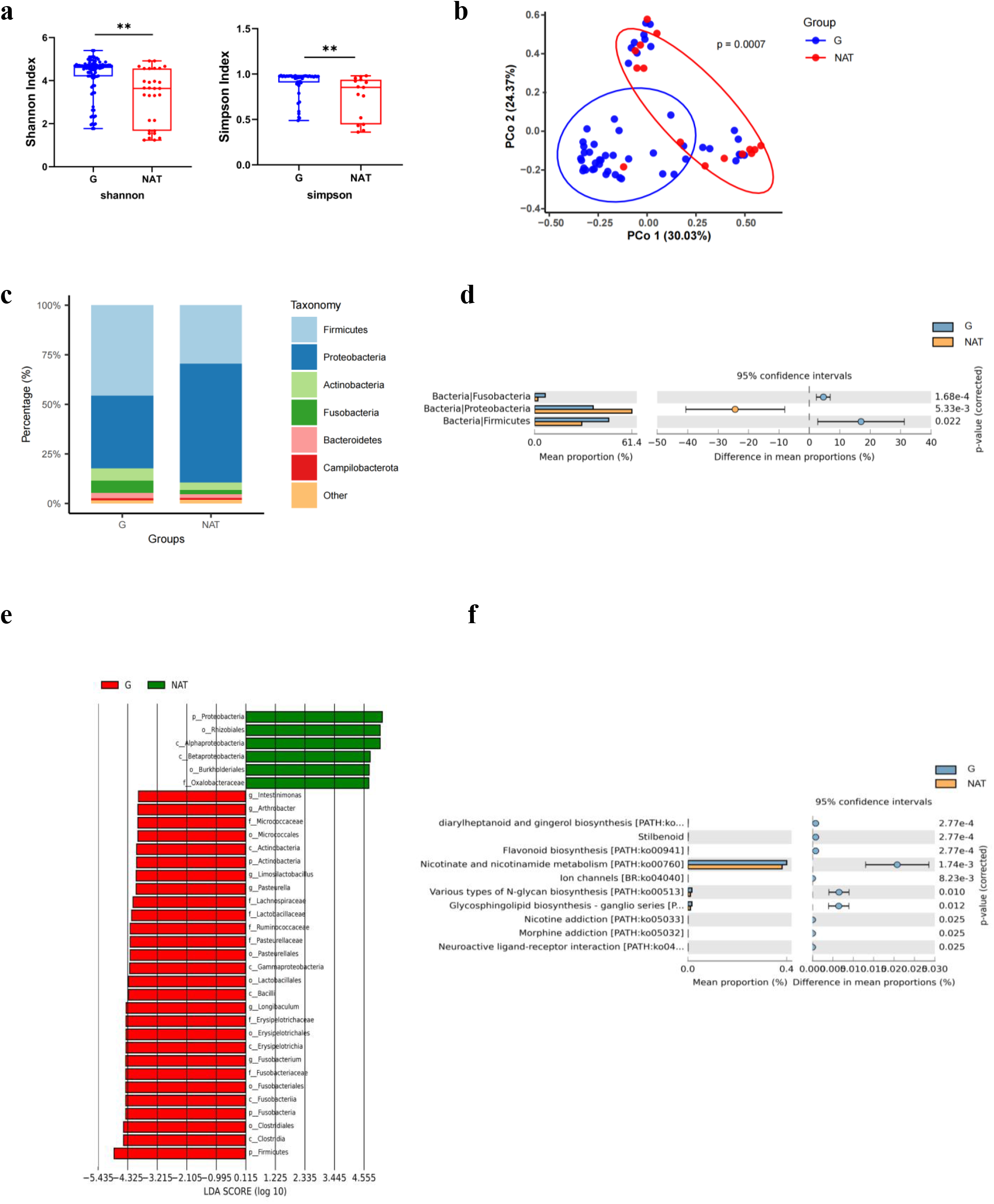
Profiling of tumor associated microbiota in human glioma tissue. **a** Species diversity differences between the G and NAT groups were estimated by the shannon, and Simpson indices. \**P* < 0.05; \*\**P*< 0.01; NS, not significant. **b** PCoA was shown along the first two principal coordinates of Bray-Curtis distances for G and NAT. The P value was calculated by PERMANOVA. G group (blue dots); NAT group (red dots), where dots represent individual samples. **c** Microbiome community structure at the phylum levels compared in G and NAT. **d** Welch’s t-test results for evaluating the relative abundance of significantly different microbiota at the phylum level. G (blue) and NAT (yellow) groups for bars and dots. **e** The distribution bar diagram based on the LEfSe analysis (LDA score (log 10)>4) in G and NAT. **f** Enrichment pathways for prediction of microbial function between G and NAT groups based on PICRUSt analysis. G, glioma tissue; NAT, adjacent normal brain tissue.

We performed linear discriminant effect size analysis (LEfSe) to identify potential glioma biomarkers in the intratumoral microbiota. We found 50 discriminatory OTUs as key discriminants, including six genera such as *Fusobacterium, Longibaculum*, *Intestinimonas*, *Pasteurella*, *Limosilactobacillus* and *Arthrobacter* (all with LDA scores (log10) > 4) (Fig. 2e). These genera were significantly enriched in the G group.

To characterize functional alterations in intratumoral bacteria, we used PICRUSt to predict functional orthologs between the G and NAT groups based on the Kyoto Encyclopedia of Genes and Genomes (KEGG). Following Bonferroni correction, we found 10 pathways enriched between G group and NAT group, including neuroactive ligand-receptor interaction (Fig. 2f).

To assess potential contributing factors to microbial diversity, we conducted stratification analysis by clinical features, including age, sex, tumor size, WHO grade and Ki-67. Interestingly, no significant relationship was observed between alpha diversity and these clinical characteristics (Additional file 3: Fig. S2a-e); however, beta diversity was found to be significantly associated with WHO grade of glioma and Ki-67 expression (Additional file 3: Fig. S2f, g).

Additionally, to explore the relationship between intratumoral microbiota and gut microbiota, we collected fecal samples from the same cohort of patients and performed 16S rRNA sequencing. Strikingly, we discovered significant differences in β diversity between tumor tissue and fecal samples from glioma patient. Conversely, the differences in α diversity were not found to be significant (Additional file 3: Fig. S3a, b).

### Morphological characteristics of bacteria in human glioma tissues

To further validate the existence of bacteria in glioma tissues, we employed serial sections for immunohistochemical staining of bacterial lipoteichoic acid (LTA) and lipopolysaccharide (LPS), as well as FISH staining for 16S rRNA. Remarkably, we observed the presence of bacterial LPS and RNA signals at the same location, whereas LTA signal was not detected (Fig. 3a). Subsequently, to determine the localization of bacteria in glioma tissue, we quantified the cells co-expressing GFAP and LPS, as well as CD68 and LPS, using multicolor immunofluorescence staining on tissue sections. Statistical analysis revealed a significantly higher abundance of GFAP^+^LPS^+^ cells compared to CD68^+^LPS^+^ cells in glioma tissues (Fig. 3b), This observation suggests that bacterial LPS is more prevalent in tumor cells than macrophages. Additionally, we performed simultaneous multicolor immunofluorescence on adjacent normal brain tissue, revealing a lower bacterial LPS signal compared to glioma tissue (Additional file 3: Fig. S4a, b).

**Fig. 3.**
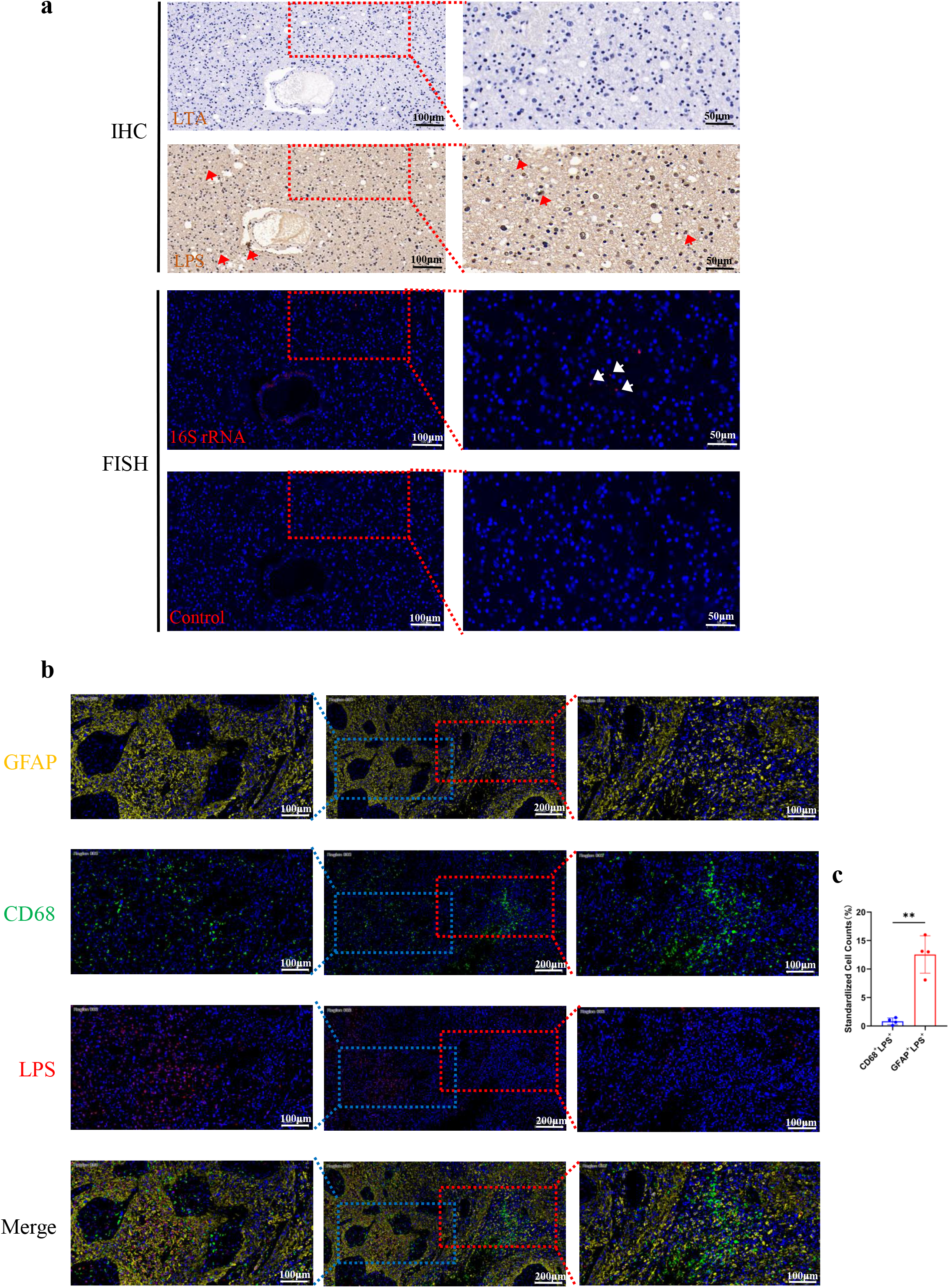
Morphological characteristics of bacteria in human glioma tissue. **a** Serial paraffin sections of human glioma samples were stained for bacterial LPS and LTA immunohistochemical staining, and bacterial 16S rRNA FISH staining, where the Control group used a negative probe without 16S rRNA. The left column is the image taken by 20x lens, scale bar = 100 μm. The right column is the 40x magnified image of the framed area, scale bar = 50 μm. LPS expression and 16S rRNA FISH signal are positive at the locations marked by red and white arrows, respectively. **b** Human glioma tissue samples were stained with multicolor immunofluorescence. GFAP (yellow), CD68 (green), LPS (red) and DAPI (blue) labeled tumor cells, macrophages, bacteria, and nuclei, respectively. The middle column displays 10x images with scale bar of 200 μm. The leftmost and rightmost columns show 20x images of the tumor cell and macrophage cluster areas, respectively, framed in the 10x images. Scale bar for 20x images is 100 μm. **c** Statistics results of cell counts of GFAP+LPS double-positive cells and CD68+LPS double-positive cells after immunofluorescence staining experiments followed by panoramic scanning. n=4; \*\**P* < 0.01.

### Profiling of microbiota-associated host gene expression in human glioma tissues

To understand the potential interactions between intratumoral microbiota and host differential gene expression, we conducted transcriptome sequencing on the same set of samples and assessed their associations using Spearman correlation analysis. Firstly, principal component analysis (PCA) revealed a significant separation between G and NAT group (Additional file 3: Fig. S5a). Furthermore, we observed remarkable alterations in the abundance of various mRNAs within glioma tissues compared to adjacent normal brain tissues (Additional file 3: Fig. S5b). According to our definition, we identified 594 differentially expressed genes (Additional file 4). The KEGG pathway enrichment analysis was performed to analyze these two groups of differentially expressed genes. Additional file 3: Fig. S5c shows the top 30 enriched terms, including neuroactive ligand-receptor interaction and glioma.

Next, we performed spearman analysis to investigate the potential relationships between all the differentially expressed genes and alpha diversity. Intriguingly, we identified 52 differentially expressed genes that exhibited significant correlations with alpha diversity (Fig. 4a). Building upon these findings, we further explored the association between these 52 differentially expressed genes and six differential bacteria. Notably, our results revealed that 28 of these differentially expressed genes displayed a predominantly negative correlation with differential bacteria (Fig. 4b). Importantly, these 28 differentially expressed genes belong to the downregulated genes in glioma, and KEGG pathway enrichment analysis demonstrated their enrichment in pathways such as cholinergic synapse, serotonergic synapse, glutamatergic synapse,and dopaminergic synapse (Fig. 4c).

**Fig. 4.**
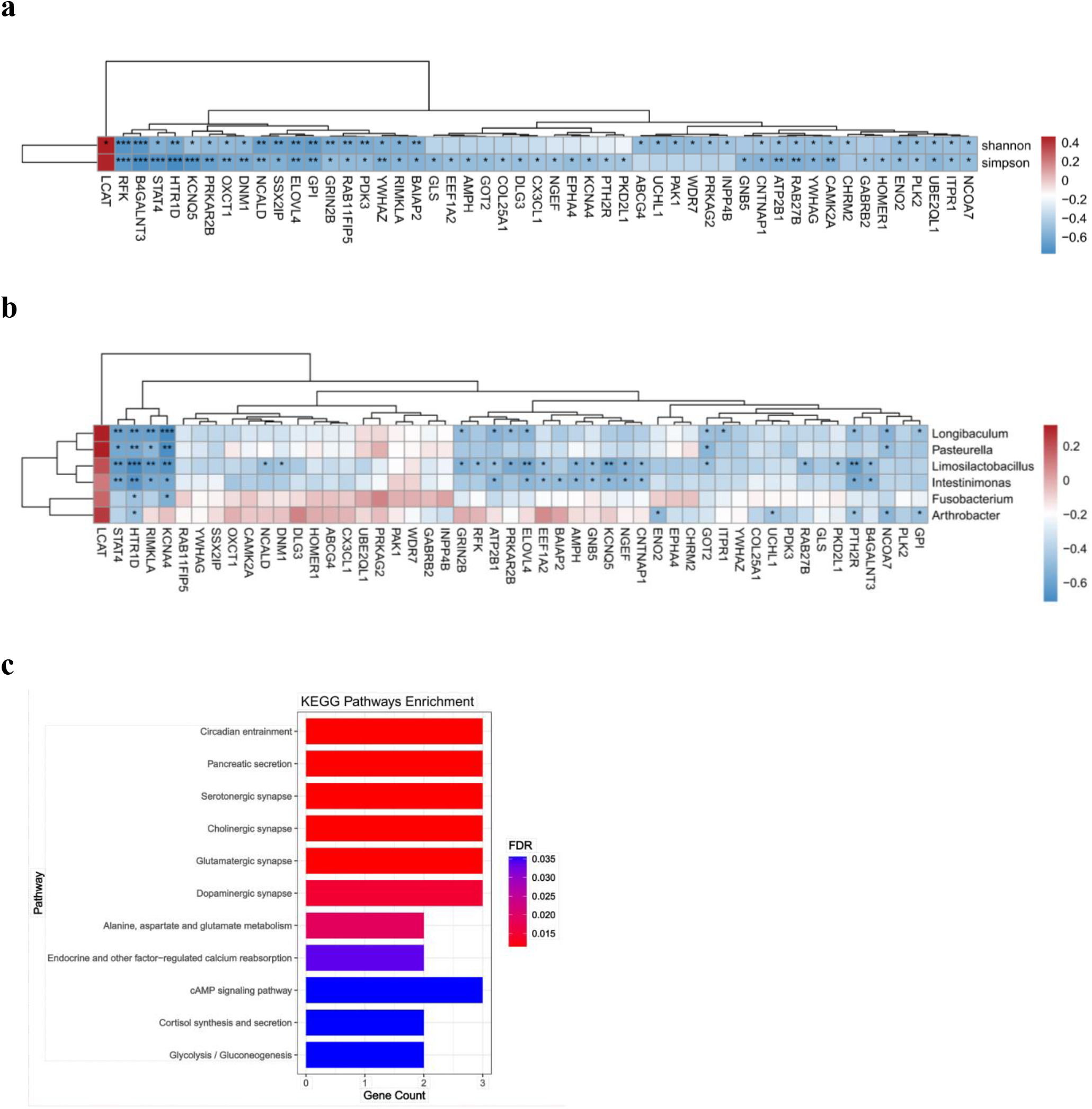
Gene expression associated with intratumoral microbiota in human glioma tissue. **a** Heatmap of Spearman correlation analysis between intratumoral microbiota alpha diversity and host gene expression. **b** Heatmap of Spearman correlation analysis between six differential bacteria abundance and host gene expression, red and blue indicate positive and negative correlations, respectively. \**P* < 0.05; \*\**P* < 0.01; \*\*\**P* < 0.001. **c** Bar diagram of KEGG pathway enrichment analysis of 28 differentially expressed genes associated with six differential bacteria abundance.

### Profiling of microbiota-associated metabolites in human glioma tissues

We conducted metabolomic assays on the same set of glioma tissues and identified microbe-associated metabolites with microbiomic data. Remarkably, the PCA plots demonstrated a clear separation of the metabolome between G and NAT group, both in ES + and ES - (Additional file 3: Fig. S6a, b). Furthermore, the OPLS-DA score scatterplots exhibited superior disjunction between the G and NAT group, further confirming the distinct metabolic profiles of glioma tissues compared to adjacent normal tissue controls, in both ES + and ES - (Fig. 5a, b).

**Fig. 5.**
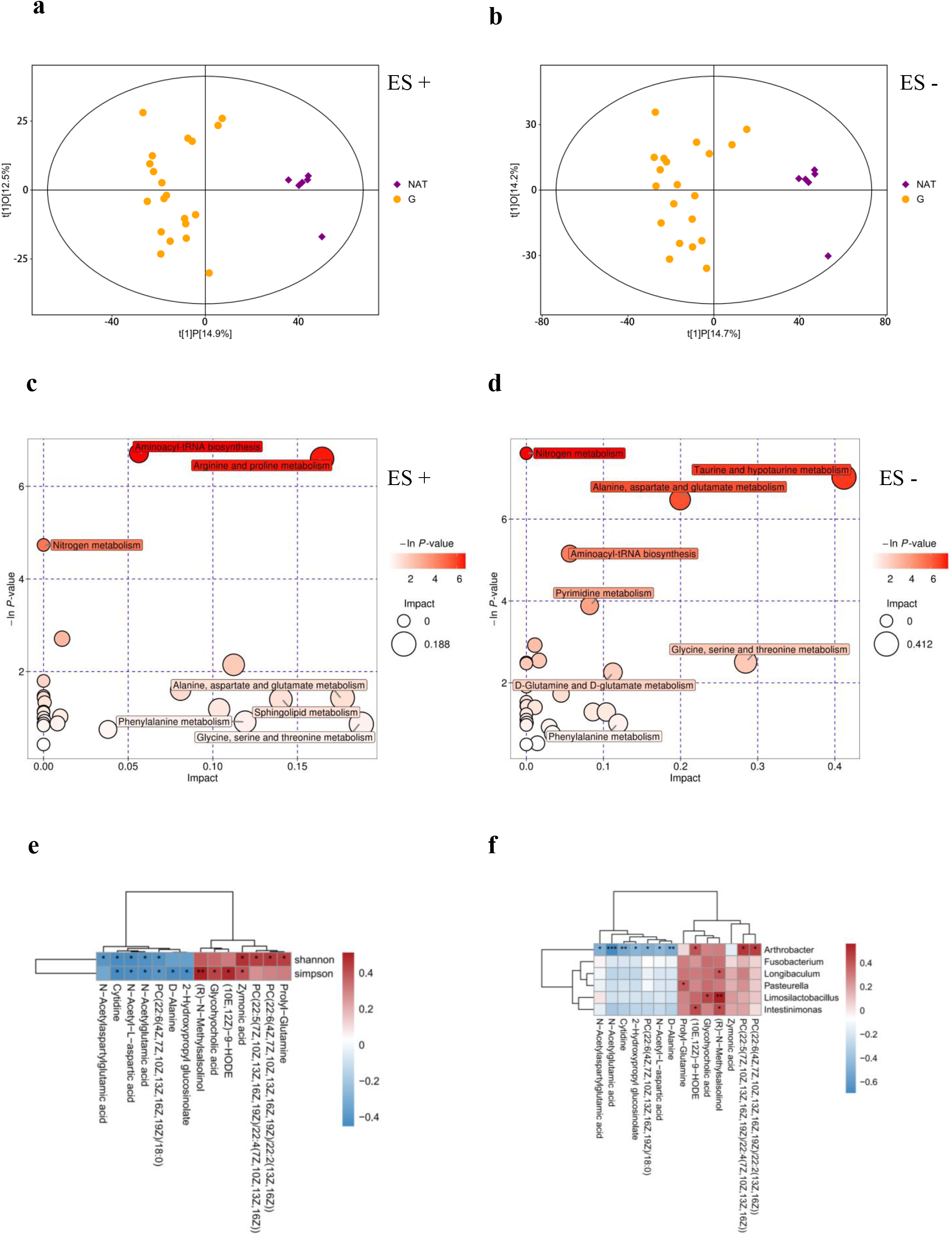
Profiling of tumor-associated metabolites in human glioma tissue. **a-b** Score scatter plot of OPLS-DA model between G and NAT group. **c-d** Metabolic pathway analysis of differential metabolites between G and NAT group. **e** Heatmap of Spearman correlation analysis between the differential metabolites and alpha diversity. **f** Heatmap of Spearman correlation analysis between the differential bacteria and metabolites, red and blue indicate positive and negative correlations, respectively. \**P* < 0.05; \*\**P* < 0.01; \*\**P* < 0.001

To identify differential metabolites, the first principal component of variable importance in the projection (VIP) of the OPLS-DA model was obtained. We identified metabolites with VIP >1 and *P* < 0.05 as differential metabolites. We identified 79 differential metabolites in ES + and 44 differential metabolites in ES - (Additional file 5), which were visualized with a hierarchical clustering heatmap (Additional file 3: Fig. S6c, d). Subsequently, all the differential metabolites underwent regulatory pathway analysis to identify the metabolic pathways that were highly correlated with the metabolites. Our analysis revealed six significantly abnormal metabolic pathways in G group, including aminoacyl-tRNA biosynthesis, arginine and proline metabolism, nitrogen metabolism, taurine and hypotaurine metabolism, alanine, aspartate and glutamate metabolism and pyrimidine metabolism. (Fig. 5c, d).

To further unravel the metabolites associated with the intratumoral microbiota in glioma, we performed spearman analysis between all the differential metabolites and alpha diversity. Remarkably, we identified 16 metabolites that exhibited correlations with alpha diversity (Fig. 5e). Subsequently, we explored the associations between these 16 differential metabolites and six differential bacteria. Our results showed some interesting correlations: (R)-N-methylsalsolinol displayed positive correlations with *Longibaculum*, *Limosilactobacillus* and *Intestinimonas*; while N-acetylglutamic acid, N-acetyl-L-aspartic acid, N-acetylaspartylglutamic acid and D-alanine exhibited negative correlations with *Arthrobacter* (Fig. 5f). It is worth noting that (R)-N-methylsalsolinol, a dopaminergic neurotoxin, has been found to increase in cerebrospinal fluid of Parkinson’s disease.[12] N-acetylaspartylglutamic acid (NAAG) and N-acetyl-L-aspartic acid (NAA), known as neurotransmitters and their precursor substances, have been reported to inhibit the differentiation of glioma stem cells.[13] Additionally, D-Alanine, a peptidoglycan constituent in bacterial cell walls, has implications as a biomarker and treatment for schizophrenia.[14]

### Integrated multi-omics analysis of human glioma tissues

As demonstrated in the previous sections, we have identified significant correlations between multiple genes, metabolites and the microbiota within glioma. In this section, we further explored the complex interactions among these three factors. Network analysis unveiled a meaningful correlation, forming interconnected networks between intratumoral microbiota, tissue metabolites and host genes (Fig. 6a and c). To assess whether genes mediate microbial effects on tumor metabolism, we performed a mediation analysis. Remarkably, we found that HTR1D and STAT4 are associated with a majority of the characteristic bacteria and differential metabolites (Fig. 6b). However, the mediation effects of the 10 pathways mediated by HTR1D and STAT4 were not statistically significant (P_mediation_ > 0.05) (Additional file 3: Fig. S7a, b). Further investigation focused on evaluating the function of metabolites in mediating the impact of microbiota on host gene expression. Our results indicated that N-acetylglutamic-acid, PC(22:5(7Z,10Z,13Z,16Z,19Z)/22:4(7Z,10Z,13Z,16Z)) and (R)-N-methylsalsolinol displayed correlations with some characteristic bacteria and some differentially expressed genes (Fig. 6d). Notably, mediation analysis showed that *Arthrobacter* causally contributed to RFK through N-acetylglutamic-acid (P_mediation_ = 0.02) (Fig. 6e). Additionally, *Longibaculum* causally contributed to GRIN2B through PC(22:5(7Z,10Z,13Z,16Z,19Z)/22:4(7Z,10Z,13Z,16Z)) (P_mediation_ = 0.016) (Fig. 6f).

**Fig. 6.**
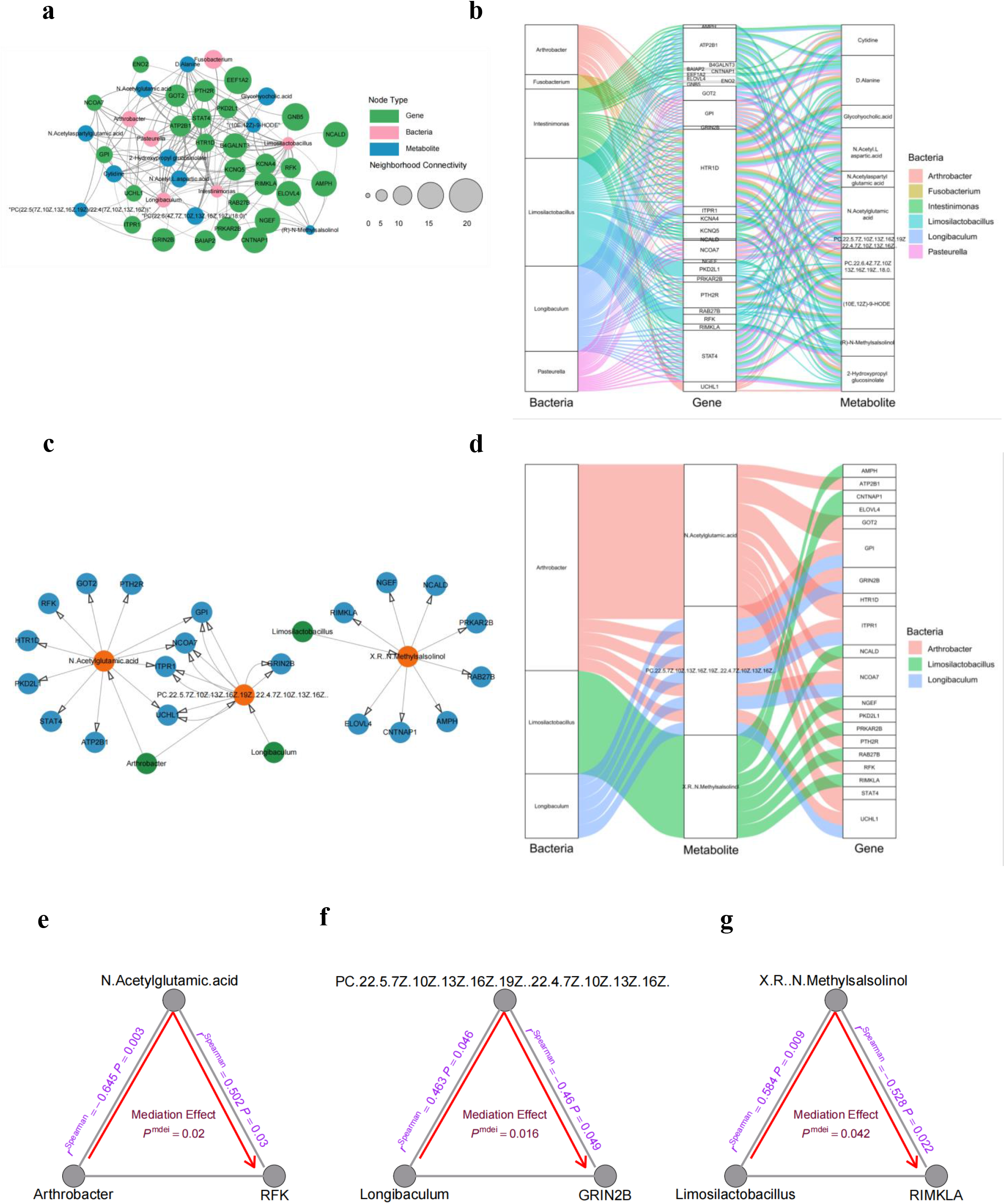
Integrated multi-omics analysis of human glioma tissues. **a** Network mapping of all differential metabolites and differential bacteria associated with differentially expressed genes in gene association analysis. **b** A Sankey diagram of multi-omics networks in glioma. **c** Network mapping of all differentially expressed genes (blue circles) and differential bacteria (green circles) associated with differential metabolite (orange circles) in metabolite association analysis. **d** A Sankey diagram of multi-omics networks in glioma. The size of each rectangle shows the degree of connectivity of each bacteria, gene, or metabolite. **e** Arthrobacter causally contributed to RFK through N-Acetylglutamic-acid (P _mediation_ = 0.02). **f** *Longibaculum* causally contributed to GRIN2B through PC(22:5(7Z,10Z,13Z,16Z,19Z)/22:4(7Z,10Z,13Z,16Z)) (P_mediation_ = 0.016). **g** *Limosilactobacillus* causally contributed to RIMKLA through (R)-N-Methylsalsolinol (P_mediation_ = 0.042). The gray lines indicate the associations among bacteria, metabolites, and genes, with corresponding r^Spearman^ values and *P*-values. The red arrowed lines indicate the bacterial effects on gene expression mediated by metabolites, with the corresponding mediation *P*-values.

Moreover, *Limosilactobacillus* causally contributed to RIMKLA through (R)-N-methylsalsolinol (P_mediation_ = 0.042) (Fig. 6g). N-acetylglutamic-acid (NAG) has been found to be involved in the regulation of NAAG degradation.[15] RFK, also known as riboflavin kinase, is the enzyme responsible for synthesizes flavin mononucleotide (FMN).[16] Evidence suggests that FMN can improve the degeneration of dopaminergic neurons.[17] PC(22:5(7Z,10Z,13Z,16Z,19Z)/22:4(7Z,10Z,13Z,16Z)) represents a phosphatidylcholine containing docosapentaenoic acid (DPA), which has been detected in the metabolomics of a variety of neurologic diseases.[18] DPA is an omega-3 polyunsaturated fatty acid with protective effects on neurons.[19] GRIN2B is a subunit of the NMDA receptor, which plays a crucial role in neural development[20], and is implicated for the communication between gut microbiota and the brain.[21] RIMKLA, also known as ribosomal modification protein rimK-like family member A, has been identified as the synthetase of NAAG.[22] In conclusion, our integrative analysis indicated that intratumoral microbiota of glioma may affect the expression of neuron-related genes through some metabolites related to neuronal function.

### Integration of the transcriptome and metabolome analysis reveals the potential role of *Fusobacterium nucleatum* in glioma mouse model

Remarkably, we detected an abundance of *Fusobacterium* in glioma tumor tissue (Fig. 2e). To further confirm the presence of *Fusobacterium* in glioma, we stained human glioma tissue and matched adjacent normal tissue samples using Cy3-labeled *Fusobacterium* probe. As shown by the results, *Fusobacterium* levels in the tumor samples were higher than those in the matched normal brain tissue (Additional file 3: Fig. S8a, b). Given that Fn accelerates the development of colorectal[23] and breast[24] cancers, we used the subcutaneous glioma xenograft mouse model to examine whether it also affects tumor progression in glioma, following the scheme in Fig. 7a. Briefly, mice were divided into four groups and performed intratumoral injections of PBS, Fn, MTZ, or Fn combined with MTZ. Fig. 7b shows that Fn group had a much larger tumor weight than the PBS and Fn+MTZ groups. Consistently, the trend persisted when evaluating tumor size (Fig. 7c). These results suggest that Fn accelerates tumor growth, while Fn-induced tumor exacerbation can be prevented by metronidazole treatment.

**Fig. 7.**
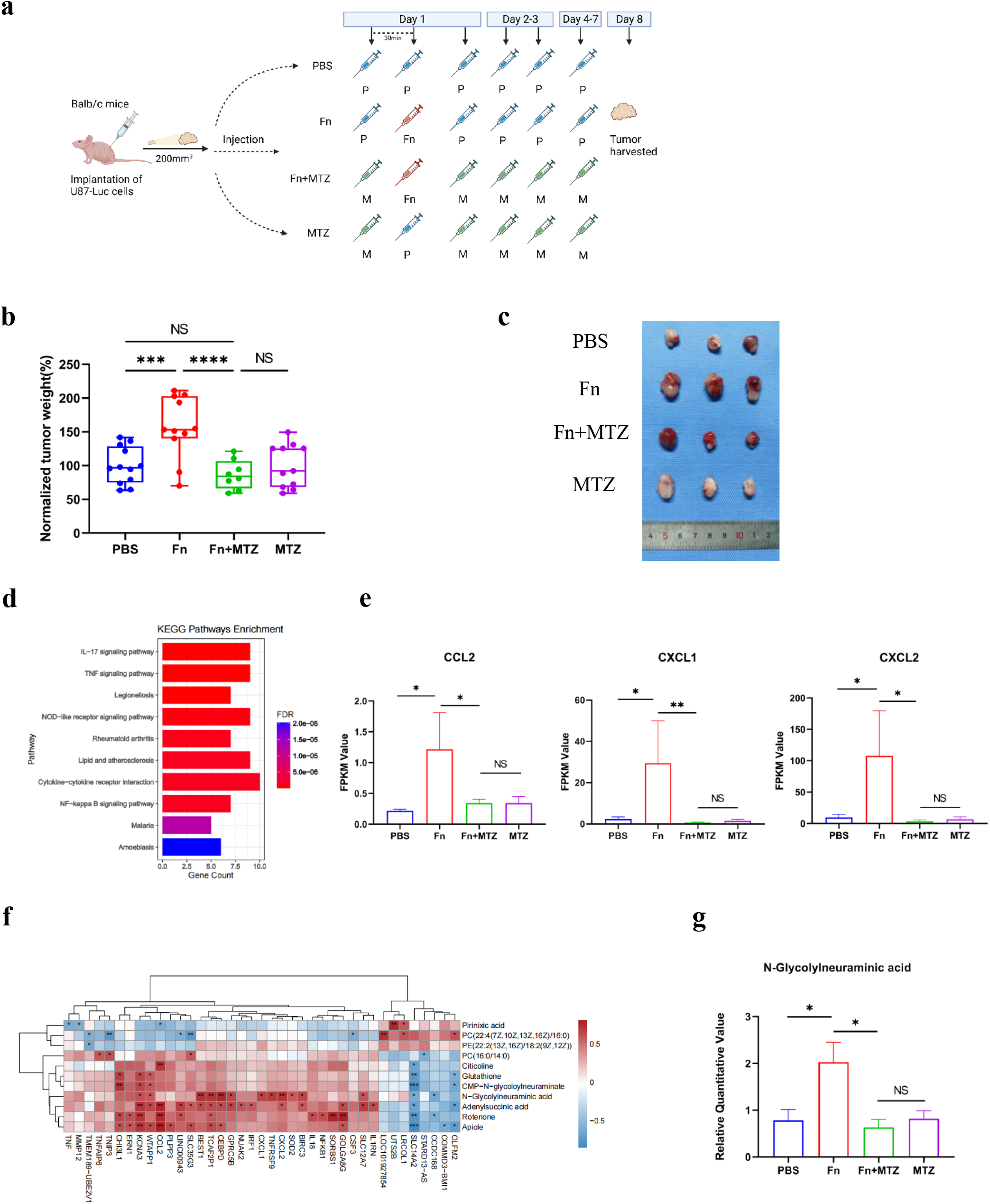
Fusobacterium nucleatum accelerates tumor growth in glioma mouse model. **a** Schematic fig. of generating subcutaneous glioma xenograft mouse model, the mice receiving intratumoral injections of PBS, Fusobacterium nucleatum (Fn), Fn along with metronidazole (MTZ) treatment or metronidazole and tumor harvested. **b** Tumor weights were normalized as percentages relative to the average tumor weight in the PBS group for each experiment (set as 100% tumor weight). Each symbol represents one mouse. **c** Representative tumors post-harvest. **d** The top 10 Bar diagram of KEGG pathway enrichment analysis of differentially expressed genes in tumor tissues from the PBS group, Fn group and Fn+MTZ group. **e** FKPM value of CCL2, CXCL1 and CXCL2 in the PBS, Fn, Fn+MTZ, and MTZ groups following transcriptomics analysis of respective tumor tissues. **f** Heatmap of Spearman correlation analysis between differential metabolites and some differentially expressed genes with obvious correlation. **g** Relative quantitative value of N-glycolylneuraminic acid in the PBS, Fn, Fn+MTZ, and MTZ groups following metabolomic analysis of respective tumor tissues. \*\*\*\**P* < 0.0001; \*\*\**P* < 0.001\*\**P*<0.01; \**P* <0.05; NS, not significant. n=3

To gain deeper insights into the specific mechanism underlying Fn promotion of glioma growth, we collected tumor tissues from the four mice groups and performed a transcriptomic analysis. The differentially expressed genes among these groups are detailed in Additional file 6. Venn diagram revealed that there were 70 genes overlapping between Fn vs PBS group and Fn vs Fn+MTZ group, which were not in Fn+MTZ vs MTZ group (Additional file 3: Fig. S9a). Subsequently, we performed KEGG pathway enrichment analysis on these 70 genes, leading to the identification of significant gathering of IL-17 signaling pathway and TNF signaling pathway (Fig. 7d). Furthermore, this analysis encompassed critical genes such as CCL2, CXCL1 and CXCL2. Fig. 7e visually demonstrates the expression levels of CCL2, CXCL1 and CXCL2 of the Fn group were markedly higher compared to those in the other three groups. Notably, previous studies have revealed the close association of these genes with Fn in promoting tumor progression.[25–27]

Furthermore, we conducted metabolomic analyses on four groups of mouse tumor tissues that received distinct treatments. The differential metabolites among these groups are detailed in Additional file 7. Venn diagram revealed the presence of 11 metabolites that overlapped between Fn vs PBS group and Fn vs Fn+MTZ group, but were not observed in Fn+MTZ vs MTZ group (Additional file 3: Fig. S9b). Subsequently, we performed correlation analysis between these 11 differential metabolites and the 70 differentially expressed genes. Notably, N-glycolylneuraminic acid exhibited a strong correlation with numerous differentially expressed genes (Fig. 7f). Furthermore, Fig. 7g visually illustrates that the expression of N-glycolylneuraminic acid of the Fn group were remarkably higher compared to those in the other three groups. Remarkably, as a sialic acid, N-glycolylneuraminic acid (Neu5Gc) is regarded as a potential human cancer biomarker.[28] Previous studies have reported the abundance of Neu5Gc in mouse brain tumor tissues.[29] Together, these data suggested that Fn promoted glioma growth by increasing the levels of N-acetylneuraminic acid and the expression of CCL2, CXCL1, and CXCL2.

## Discussion

With the growing attention to the gut-brain axis[30], the gut microbiota has been found to be involved in the development, progression and treatment of glioma by metabolically modulating the epigenetic[31] and immune microenvironments[32]. Recently, emerging evidence has also shown a potential function of the intratumoral microbiome in tumor behavior[33] and treatment responses[34]. It is of great importance for cancer therapy to elucidate the molecular features of distinct tumor subtypes and predict clinical prognosis from a microbiome perspective. However, the presence of intratumoral microbiota has been questioned due to their low abundance and the possibility of contamination. In 2020, Nejman et al.[9] performed a comprehensive bacterial detection on seven types of solid tumors and proposed a guideline for handling contamination in low-biomass samples. In our investigation, to obtain more reliable data, we also followed this guideline and performed contaminant filtering. By analyzing 16S rRNA gene sequencing data, we identified six differential bacteria in glioma tissues compared to matched relatively normal brain tissues, including *Fusobacterium*, *Longibaculum*, *Intestinimonas*, *Pasteurella*, *Limosilactobacillus* and *Arthrobacter*. It is well known that tumor hypoxia, a centerpiece of disease progression mechanisms, is also a common pathological feature in glioma. We found that most of the genera enriched in glioma tissue were anaerobic, which does not seem to be a coincidence. In fact, the high degree of hypoxia in the tumor, immunosuppressive microenvironment, and disturbed vascular system are favorable conditions for rapid bacterial colonization, growth, and replication within the tumor.[35]

Furthermore, we observed a marked discrepancy between the microbial composition of fecal samples and tumor samples from the same cohort of glioma patients. This implies that the gut microbiota is not the only possible source of intratumoral bacteria in glioma. These bacteria may also originate from the oral cavity, or adjacent brain tissue. We make the following speculations: 1) glioma might change the local microenvironment, such as blood–brain barrier disruption and immunosuppression, which enables bacteria to infiltrate the tumor through the hematogenous or the neuronal retrograde pathways. 2) These bacteria could have been present in the brain tissue prior to tumorigenesis, and those that adapted to the tumor microenvironment survive and grow during tumor development. Of course, these speculations should be investigated in future studies.

*Fusobacterium* is a genus of obligate anaerobic rods, and *Fusobacterium nucleatum* has been reported as a bacterium closely related to tumorigenesis.[36] *Fusobacterium* was found to be enriched in stool samples from glioma patients.[37] Here, it was found to be enriched in the tumor tissues of glioma patients. *Longibaculum*, belonging to the bacterial genus within the family *Erysipelotrichaceae*, has not been extensively investigated in the context of glioma. However, studies have indicated its involvement in weight-independent improvement of blood glucose subsequent to gastric bypass surgery.[38] In a mouse model of glioma, *Intestinimonas* was reported to exhibit a continuous increase in abundance during the progression of tumor growth in a mouse model of glioma.[39] Remarkably, *Intestinimonas* possesses distinctive metabolic capabilities that allow it to produce butyrate from both carbohydrates and amino acids.[40] *Pasteurella* is a genus of opportunistic pathogens, and one of its species, *Pasteurella multocida*, has been reported to cause bacterial meningitis.[41] Although investigations thus far have not illuminated any direct link between *Pasteurella* and glioma, the presence of Pasteurella multocida toxin (PMT), characterized by profound mitogenic activity and carcinogenic potential[42], presents an intriguing possibility. It is conceivable that PMT may serve as a pivotal mediator of the association between *Pasteurella* and glioma pathogenesis. *Limosilactobacillus reuteri*, extensively studied for its protective properties[43], has recently been implicated in the induction of multiple sclerosis.[44] *Arthrobacter* strain NOR5 has exhibited remarkable proficiency in facilitating the complete degradation of nornicotine[45], a direct precursor of tobacco-specific nitrosamines (TSNAs) known for their potent carcinogenic properties.[46] Furthermore, *Arthrobacter citreus* strains can metabolize caprolactam, thereby generating glutamate[47]-an important excitatory neurotransmitter in the central nervous system. Based on these findings, it becomes conceivable that Arthrobacter holds probiotic potential. Notably, our investigation also revealed an association between Arthrobacter and NAA as well as NAAG.

It is undeniable that culturing bacteria from fresh glioma tissues is the critical evidence for their presence. Regrettably, our endeavors in bacterial culture encountered setbacks. Our efforts resulted solely in the cultivation of *Pseudomonas stutzeri*, a bacterium that has exhibited resistance against multiple antibiotics (data not shown). We speculate that this outcome may be attributed to the constraints posed by current medical practices, particularly the widespread utilization of prophylactic antibiotics.

Furthermore, a noteworthy observation in our study was the notable elevation of LPS signals in glioma tissue in stark contrast to adjacent normal brain tissue. LPS, a prevalent endotoxin, displays the capacity to interact with TLR4 in vivo, provoking the activation of monocytes and macrophages, thereby instigating the synthesize and subsequent release of various cytokines and inflammatory mediators.[48] Remarkably, investigations have illustrated that glioblastoma clinical samples exhibit heightened expression levels of TLR4[49], corroborating the findings of our study. Moreover, we also found that LPS signals were more prevalent in tumor cell-enriched regions, while macrophage-enriched regions had fewer signals. We propose that this may result from LPS-induced macrophage activation and subsequent phagocytic clearance by macrophages. Notably, alongside bacterial LPS, RNA, and DNA found in human gliomas, recent studies have unraveled the presence of bacterial-derived peptides on HLA molecules in glioblastoma.[50] These peptides elicit strong responses from tumor-infiltrating lymphocytes and peripheral blood memory cells, suggesting that microbial peptides activate tumor-infiltrating lymphocytes in glioblastoma.

In addition, we performed transcriptomic sequencing on glioma tissues derived from patients simultaneously. We found that 28 differentially expressed genes associated with six different bacterial genera were enriched in pathways such as serotonergic synapse, cholinergic synapse, glutamatergic synapse, and dopaminergic synapse. The integrated omics analysis revealed that intratumoral microbiome may affect the expression of neuron-related genes through bacteria-associated metabolites. Previous review articles have highlighted the ability of gliomas to exploit normal mechanisms of neuronal development and plasticity, leading to the formation of neuron–glia synapses, with subsequent enhancement of glioma proliferation.[51] Building upon these limited but intriguing pieces of evidence, we speculate that the intratumoral bacteria in glioma may be involved in the formation of neuron-glia synapses.

Additionally, a noteworthy discovery emerged as we unraveled the association of HTR1D and STAT4 with the characteristic bacteria and the majority of the differential metabolites. HTR1D, a type of 5-hydroxytryptamine (5-HT) receptor, activates intracellular signaling pathways through G proteins upon binding to serotonin.[52] Such signal transduction represents a fundamental process mediated by the monoamine neurotransmitter 5-HT. Consequently, HTR1D may be an important molecule in the neuron–glia synapses involved by intratumoral bacteria of glioma. Concurrently, signal transducer and activator of transcription 4 (STAT4) operates as a transcription factor. Prior studies employing the Oncomine database scrutinized the mRNA expression levels of STAT gene family members in glioma, revealing diminished STAT4 mRNA expression in comparison to that observed in normal controls[53]-an observation corroborated by our own inquiry. Of equal import, several studies have unveiled STAT4’s pivotal role in regulating neutrophil function, as demonstrated by the impact of STAT4 deficiency on neutrophil extracellular trap formation and antibacterial immunity.[54] Therefore, within the context of intratumoral bacteria-associated glioma, STAT4 emerges as an indispensable molecule in the tumor-associated immune regulation.

In a mouse model of glioma, *Fusobacterium nucleatum* promotes glioma growth by increasing the levels of N-acetylneuraminic acid and the expression levels of CCL2, CXCL1, and CXCL2. Notably, recent study has shed light on the activation of inflammatory response-associated genes, including TNF, INFγ, and IFNα, as well as the upregulation of chemokines CCL2, CXCL1, and CXCL10 in macrophages that phagocytose bacteria within the tumor microenvironment.[50] Encouragingly, our study aligns with these findings. Collectively, our comprehensive investigation implies a diversified landscape pertaining to the interplay between intratumoral bacteria and glioma. It is important to acknowledge the dissimilarity between the subcutaneous U87-luc tumor-bearing mouse model we employed and the original tumor microenvironment of the patients. Consequently, the detected differential genes and metabolites may also vary. We strongly advocate for future studies to harness patient-derived organoid models and mouse orthotopic tumor models to validate specific pathways experimentally-an endeavor that will constitute an aspect of our forthcoming research agenda.

Previous studies have demonstrated the ability of *Fusobacterium nucleatum* to confer resistance to chemotherapy[55] and the enhance of PD-L1 blockers[56] in colorectal cancer. Given the enrichment of Fn observed in glioma and its capability to recruit chemokines, it becomes crucial to explore the potential associations between Fn and temozolomide, a first-line chemotherapy drug for glioma, as well as the interplay between Fn and PD-L1 in glioma context. These avenues of investigation hold considerable promise for future research endeavors. The advancement of bacterial engineering technology an exciting opportunity to develop specifically modified bacteria that can serve as anti-tumor carriers. Among the six differential bacteria we have identified, there is potential for them to be utilized as target bacteria for future genetic engineering modification.

Despite the significance of our findings, our study has certain limitations. Firstly, the postulated interactions between the intratumoral bacteria and glioma necessitate validation through targeted experiments investigations. Secondly, our study sample size was relatively small, highlighting the need for multicenter, large sample studies to elucidate the relationship between key bacteria, their related metabolites and glioma prognosis. Nevertheless, our study provides novel insights into the intricate interplays between intratumoral bacteria and glioma, potentially inspiring new avenues of exploration in glioma biology. Looking ahead, an in-depth study of the intratumoral microbiota holds immense promise for advancing anti-cancer treatment.

In conclusion, a multi-omics analysis of glioma tissue showed that the intratumoral microbiome may affect the expression of neuron-related genes through bacteria-associated metabolites. In a glioma animal model, Fn increased the levels of N-glycolylneuraminic acid and the expression of genes such as CCL2, CXCL1, CXCL2.

## Acknowledgments

Thanks to Dr SongHe Guo for the kind gift of the strain ATCC25586(*Fusobacterium nucleatum*) and assistance with bacterial culture. Thanks to Dr Yan He and Dr Jiayuan Huang for carefully reading the article and giving valuable suggestions. The authors also thank all staff at the technical staff of the Clinical Biobank Centre for their kind assistance. Finally, they would also like to thank all tissue donors and their families who have kindly donated their resected specimens to the Clinical Biobank Centre. We acknowledge Biorender (©BioRender - biorender.com) since figures of this draft were made using this software.

## Author contributions

Haitao Sun conceptualized the project, designed and supervised the study. Ting Li and Zhanyi Zhao performed experiments and drafted the manuscript. Meichang Peng and Chen Wang participated execution of experiments and data analysis. Feiyang Luo polished the English language. Meiqin Zeng, Kaijian Sun and Zhencheng Fang participated in the data analysis. Yunhao Luo, Qiyuan Huang and Yugu Xie participated in the collection of samples. Jiaxuan Wang participated in analysis and interpretation of data. Jian-Dong Huang and Hongwei Zhou participated in the study supervision. All authors contributed to writing the manuscript, provided critical feedback during editing and approved the final submitted version.

## Data Availability

The raw sequence data reported in this paper have been deposited in the Genome Sequence Archive[57, 58] in National Genomics Data Center, China National Center for Bioinformation / Beijing Institute of Genomics, Chinese Academy of Sciences (GSA-Human: HRA005548) that are publicly accessible at https://ngdc.cncb.ac.cn/gsa-human. The metabolomics data reported in this paper have been deposited in the OMIX under the accession no. OMIX004935 and no. OMIX004936. (https://ngdc.cncb.ac.cn/omix).All other data supporting the findings of this study are available within the article, the supplementary information, or from the corresponding authors upon reasonable request.

## Declarations

### Ethics Approval and Consent to Participate

The study received the informed consent of patients and was approved by the local Ethics Committee (2018-SJWK-004) (2022-KY-KY-102-02). The registration has been completed on the website of the China Clinical Trial Registry Center (ChiCTR2300067789). The Animal Ethics Committee of Southern Medical University approved the animal study which was performed according to the National Institutes of Health (Approval ID: LAEC-2022-197).

## Additional material

File name: Additional file 1

File format: xlsx

Title of data: Supplementary Table 1. The clinical data of glioma patients.

File name: Additional file 2

File format: pdf

Title of data: Supplementary Methods

File name: Additional file 3

File format: pdf

Title of data: Supplementary Figure

File name: Additional file 4

File format: xlsx

Title of data: Supplementary Table 2. Differentially expressed genes in human glioma tissues.

File name: Additional file 5

File format: xlsx

Title of data: Supplementary Table 3. Differential metabolites in human glioma tissues.

File name: Additional file 6

File format: xlsx

Title of data: Supplementary Table 4. Differentially expressed genes in mouse tumor tissues.

File name: Additional file 7

File format: xlsx

Title of data: Supplementary Table 5. Differential metabolites in mouse tumor tissues.

