## Additional file 2 for "Multi-omics analysis reveals novel interplays between intratumoral bacteria and glioma"

### Supplementary Methods

#### Multiplex Immunofluorescent Assay

Endogenous peroxidase was quenched with 3% H2O2 for 30 minutes, and then blocked with blocking reagent at room temperature for 30 minutes. Primary antibody was incubated overnight in a humidified chamber at 37 °C, washed with PBS, 3x15min, and then used HRP-conjugated secondary antibody (1:500), incubated at room temperature for 30min, washed with PBS, 3x15min, and then stained with TSA-bifluorescein (no more than 60s). Then, the slides were placed in recovery/wash buffer (Abcracker) and boiled at high power in a microwave oven until the repair solution boiled, maintained for 10 seconds and turned off, and the beaker could be placed in a water bath in a basin after 5 minutes to cool to room temperature. In order, each antigen was labeled with a different fluorescent group.

#### 16S rRNA FISH

Slides were washed with 1xPBS (2x10 min, at RT) and then treated with HCl (0.2 N, 20 min) and Proteinase K (50μg/mL, 20 min) at RT. Slides were washed with 1xPBS (1x5 min, at RT) and then incubated with 200μL blocking buffer for 2 h at 55 °C. Slides were washed in PBS for 5 min and air-dried. The probe solutions (1:100 dilution, 250nM) were prepared by mixing probes with 25% hybridization buffer and hybridized for 72 h at 37 °C. Then, slides were washed in pre-warmed washing buffer (60°C) for 15 min and air-dried for 20 min. Finally, slides were mounted with 20 μL DAPI-Antifade solution for 10 min and covered with a cover slide in dark.

**DNA extraction and sequencing**

PCR amplification of the bacterial 16S rRNA gene V3-V4 regions was performed using the forward primer 341F (5'–CCTAYGGGRBGCASCAG–3') and the reverse primer 806R (5'–GGACTACNNGGGTATCTAAT–3'). Sample-specific 6-bp barcodes were incorporated into the primers for multiplex sequencing. All PCR reactions were carried out with 15 μL of Phusion® High-Fidelity PCR Master Mix (New England Biolabs), 0.2 μM of forward and reverse primers, and about 10 ng template DNA. Thermal cycling consisted of initial denaturation at 98 °C for 1 minute, followed by 30 cycles of denaturation at 98 °C for 10s, annealing at 50 °C for 30s, elongation at 72 °C for 30 s, and finally 72 °C for 5 min. PCR amplicons were purified with Qiagen Gel Extraction Kit (Qiagen, Germany) by agarose gel electrophoresis (2%). Sequencing libraries were generated using TruSeq® DNA PCR-Free Sample Preparation Kit (Illumina, USA) following the manufacturer's recommendations. The library quality was assessed on the Qubit@2.0 Fluorometer (Thermo Scientific) and Agilent Bioanalyzer 2100 system. The paired-end sequencing (2x250 bp) was performed on the Illumina NovaSeq platform (Novogene, Tianjin). Pairing and reading samples according to their unique bar codes, and cutting off the bar codes and primer sequences. We used FLASH software to merge partner readings. Quality filtering is carried out on the original reading under specific filtering conditions, and high-quality clean labels are obtained according to the formula (v0.94). EasyAmplicon (v1.12) was used to analyze downstream Amplicon bioinformatics. The full-length dereplication (derep_fulllength) command used VSEARCH (v2.15.2). Then, the Unoise3 command of USEARCH (v10.0.240) was used to denoise the non-redundant sequences into Amplicon sequence variation (ASVs). VSEARCH (v2.15.2) was also used for reference-based chimera detection. A representative sequence was selected for each ASV, and the SILVA reference database was used to annotate taxonomic information.

**mRNA-seq**

The RNA sequencing was performed on the Illumina Nova6000 sequencer (Illumina). The expression levels for each gene were normalized to fragments per kilobase of transcript per million fragments mapped to compare mRNA abundance between samples. The DEGs were submitted for GO enrichment analysis with the Bioconductor software package topGO. Significant GO terms with $P \leq 0.05$ were shown.

**Metabolomics**

25 mg of sample was weighted to an EP tube, and 500 μL extract solution (methanol: acetonitrile: water = 2: 2: 1, with isotopically-labelled internal standard mixture) was added. Then the samples were homogenized at 35 Hz for 4 min and sonicated for 5 min in ice-water bath. The homogenization and sonication cycle were repeated for 3 times. Then the samples were incubated for 1 h at -40 °C and centrifuged at 12000 rpm for 15 min at 4 °C. The resulting supernatant was transferred to a fresh glass vial for analysis. LC-MS/MS analyses were performed using an UHPLC system (Vanquish, Thermo Fisher Scientific) with a UPLC BEH Amide column (2.1 mm × 100 mm, 1.7 μm) coupled to Orbitrap Exploris 120 mass spectrometer (Orbitrap MS, Thermo). The mobile phase consisted of 25 mmol/L ammonium acetate and 25 mmol/L ammonium

hydroxide in water（pH = 9.75）(A) and acetonitrile (B). The auto-sampler temperature was 4 ℃, and the injection volume was 2 μL. The Orbitrap Exploris 120 mass spectrometer was used for its ability to acquire MS/MS spectra on information-dependent acquisition (IDA) mode in the control of the acquisition software (Xcalibur, Thermo). The raw data were converted to the mzXML format using ProteoWizard and processed with an in-house program, which was developed using R and based on XCMS, for peak detection, extraction, alignment, and integration. Then an in-house MS2 database (Biotree Biomedical Technology Corporation, Shanghai, China) was applied in metabolite annotation. The cutoff for annotation was set at 0.3.

In this experiment, SIMCA software (v16.0.2) was used to perform logarithmic transformation plus centering formatting on the data, and then automatic modeling analysis was performed to obtain the principal component analysis (PCA) model. Next, the data were log-transformed plus UV formatted using SIMCA software (v16.0.2), and the first principal component was first analyzed by OPLS-DA modeling.
