## Additional file 3 for "Multi-omics analysis reveals novel interplays between intratumoral bacteria and glioma"

### Supplementary Figure

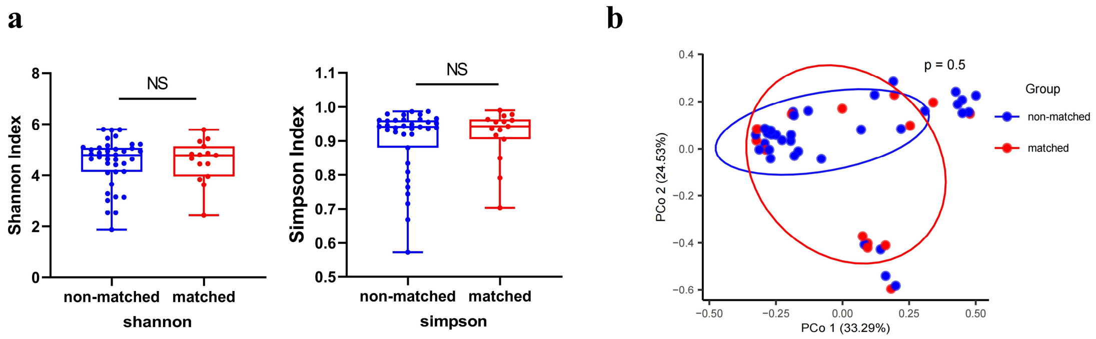


**Figure S1** Microbial diversity between non-matched and matched groups in glioma tissue.

**a** α diversity differences were estimated by the shannon, and simpson indices. NS, not significant.

**b** PCoA was shown along the first two principal coordinates of Bray-Curtis distances. The P value was calculated by PERMANOVA. non-matched group (blue dots); matched group (red dots), where dots represent individual samples.

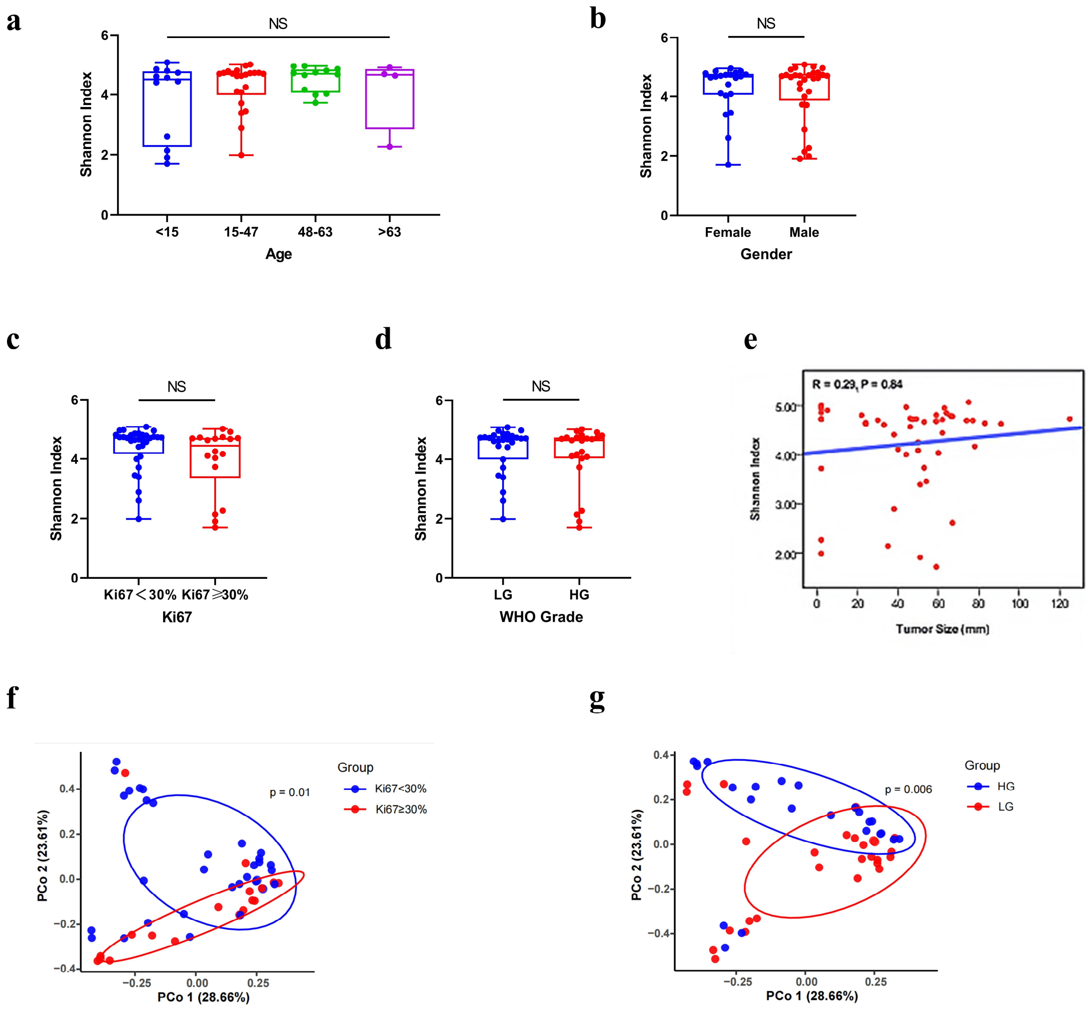


**Figure S2** Association between intratumoral microbiota and clinical features of glioma**.**

**a-e** Alpha diversity boxplots (Shannon index) in glioma patients. Analysis of following factors: Age, Gender, Ki67, WHO grade, Tumor size.

**f-g** PCoA plots of intratumoral bacteria diversity in the Ki67<30% and Ki67≥30% glioma groups and in the low-grade and high-grade glioma groups. LG, WHO I - II; HG, WHO III – IV.

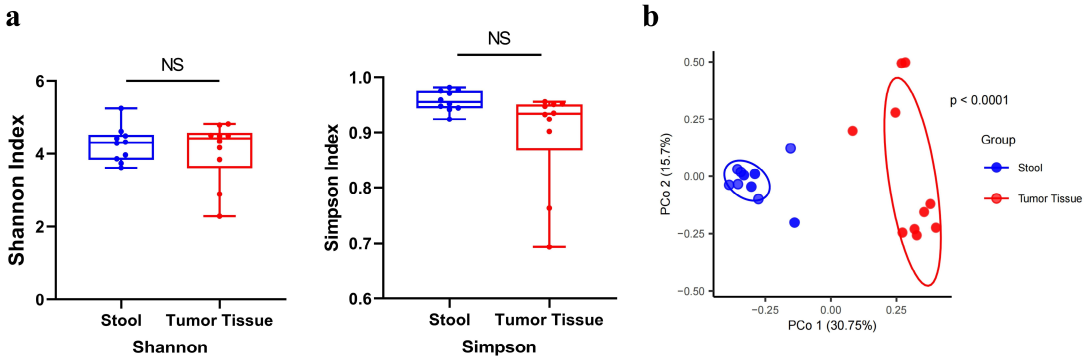


**Figure S3** Microbial diversity between stool and tumor tissue samples in glioma.

**a** α diversity differences were estimated by the shannon, and simpson indices. *$P <$ 0.05; NS, not significant.

**b** PCoA was shown along the first two principal coordinates of Bray-Curtis distances. The P value was calculated by PERMANOVA. Stool group (blue dots); Tumor Tissue group (red dots), where dots represent individual samples.

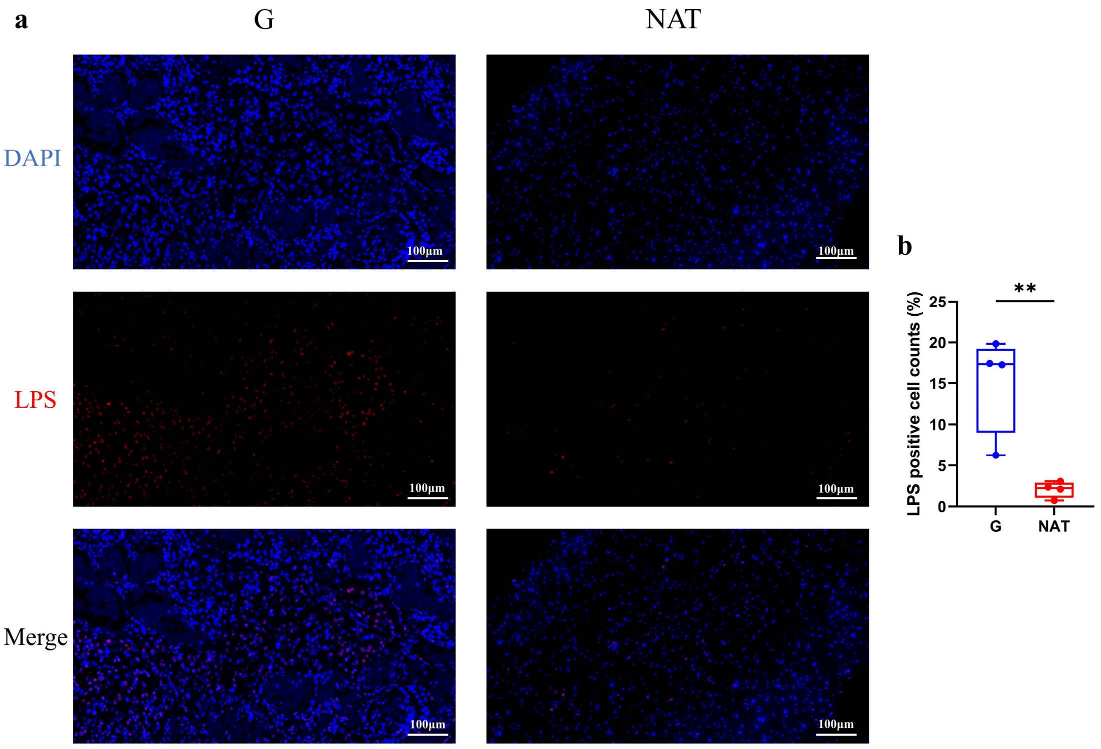


**Figure S4** Immunofluorescence staining of LPS in human glioma tissue and adjacent normal brain tissue.

**a** Immunofluorescence staining of glioma tissue and adjacent normal brain tissue with anti-LPS antibody at 20X, respectively. LPS (red) and DAPI (blue). scale bar = 100 μm.

**b** Statistics results of cell counts of LPS positive cells after immunofluorescence staining experiments followed by panoramic scanning. Each symbol represents one sample. n=4; ***P* < 0.01.

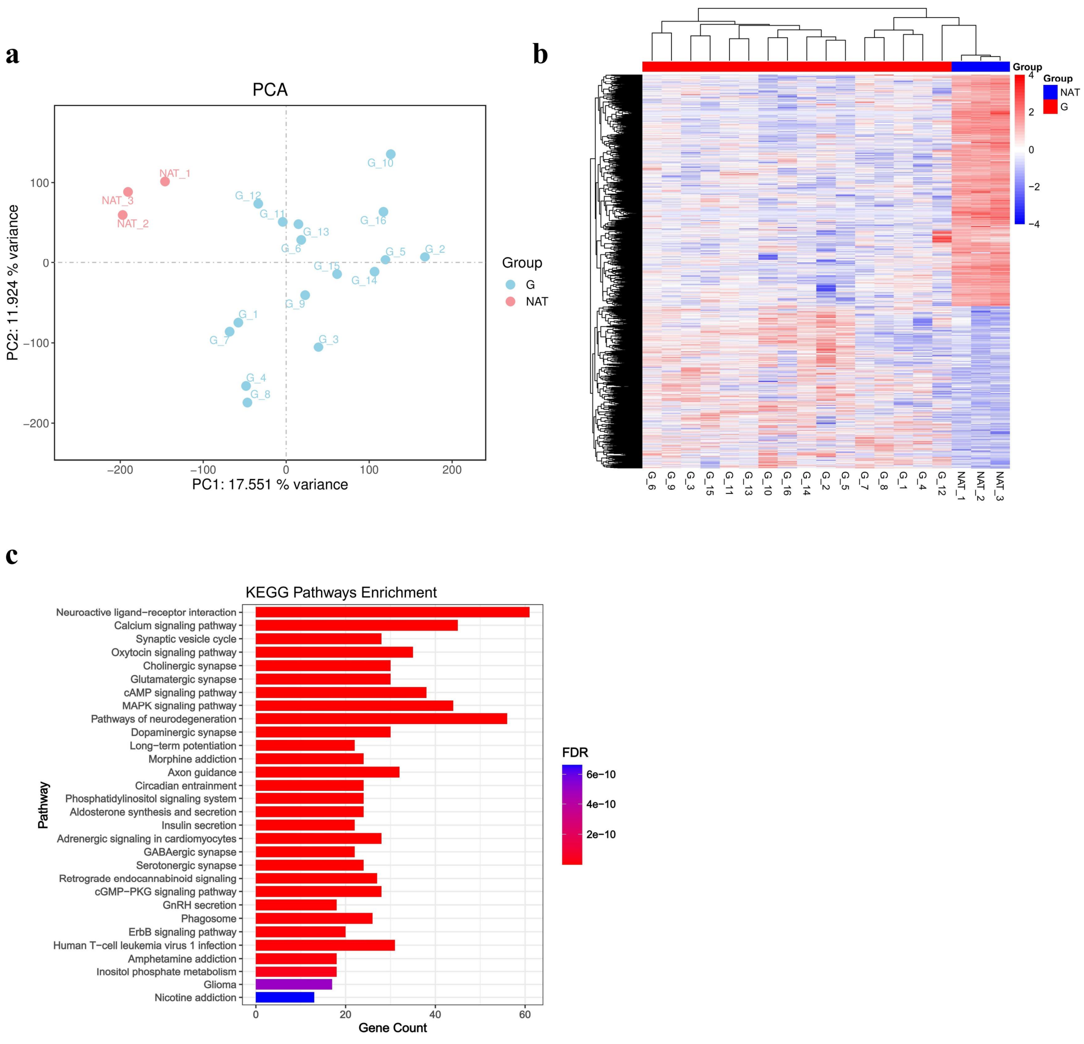


**Figure S5** Profiling of host gene expression in human glioma tissue.

**a** PCA plot of transcriptomics data collected from human glioma tissue and adjacent normal brain tissue samples.

**b** Hierarchical clustering heatmap of differential expression genes between G and NAT group.

**c** Bar diagram of KEGG pathway enrichment analysis of all differential expression genes.

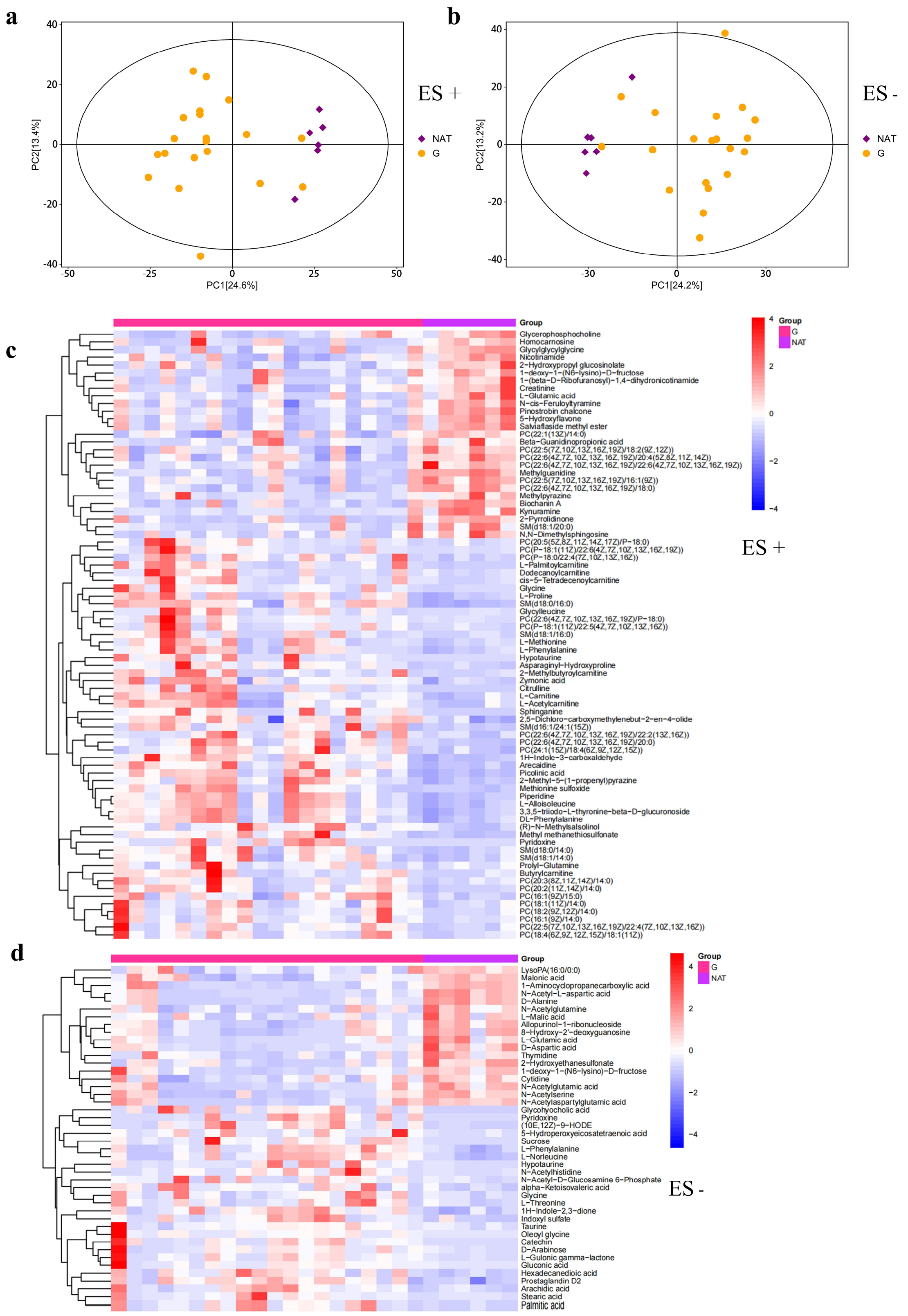


**Figure S6** Metabolomic analysis of glioma tissue and adjacent normal brain tissue.

**a-b** Score scatter plot of PCA model between G and NAT group.

**c-d** Hierarchical clustering heatmap of differential metabolites between G and NAT group.

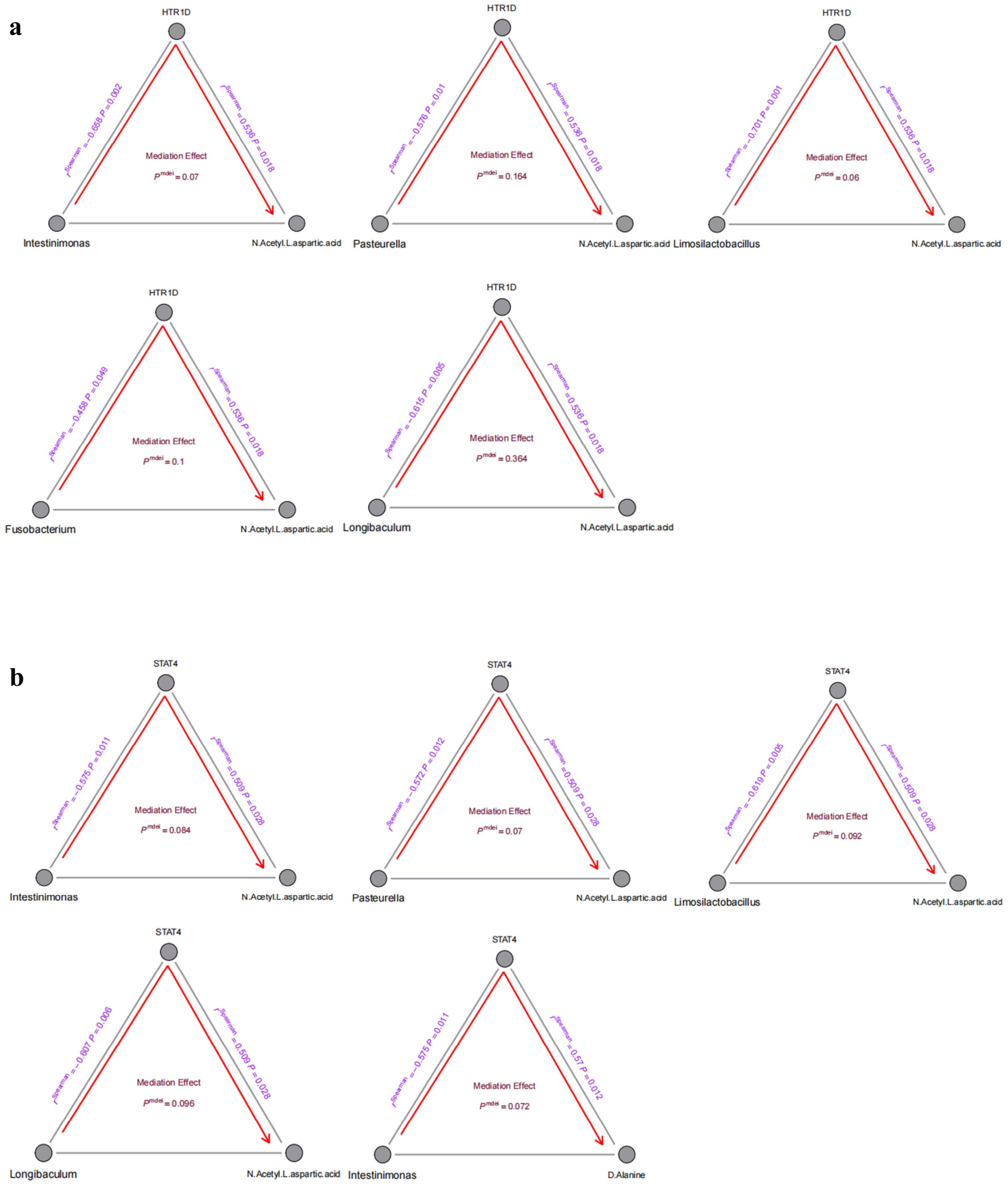


**Figure S7** Diagram of Mediation analysis

**a** Pathways mediated through HTR1D.

**b** Pathways mediated through STAT4. The gray lines indicate the associations among bacteria, metabolites, and genes, with corresponding $r^{Spearman}$ values and *P*-values. The red arrowed lines indicate the bacterial effects on gene expression mediated by metabolites, with the corresponding mediation *P*-values.

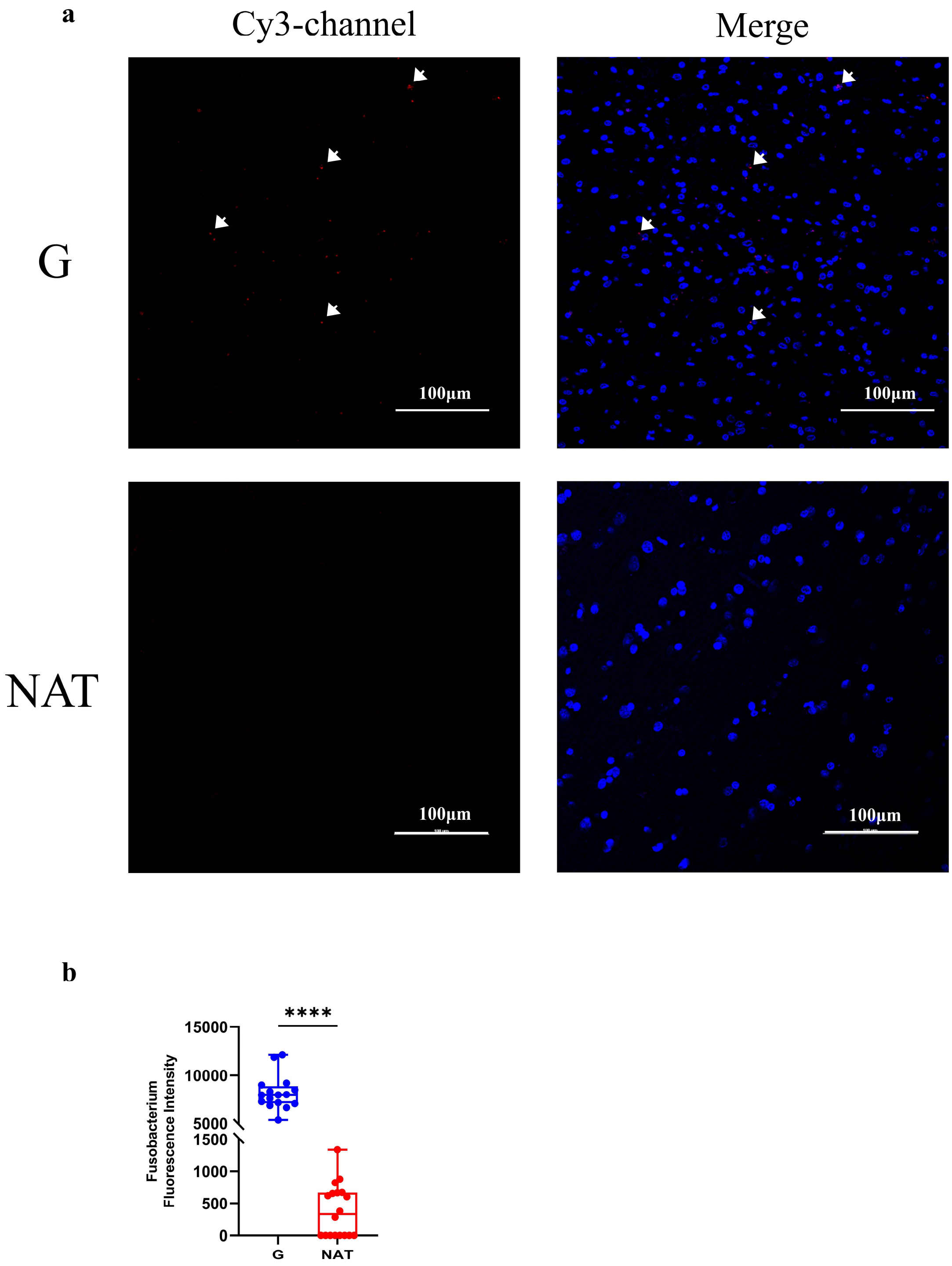


**Figure S8** FISH staining of *Fusobacterium* in human glioma tissue and adjacent normal brain tissue.

**a** FISH staining of glioma tissue and adjacent normal brain tissue with Cy3-labeled *Fusobacterium* probe at 40X, respectively. *Fusobacterium* signals (red) and DAPI (blue). scale bar = 100 μm.

**b** Statistical analysis of the levels of fluorescent signals in (A). Each dot represents one sample. G n=16; NAT n=18.; ****$P$ < 0.0001.

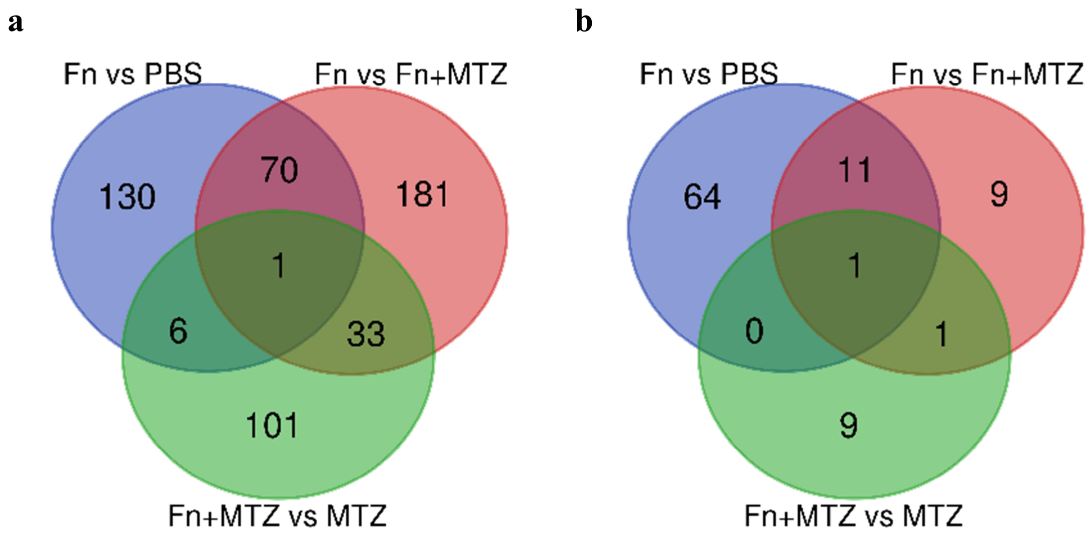


**Figure S9** Analysis of differentially expressed genes and differential metabolites in mice glioma tissues.

**a** Summary of differentially expressed genes (determined by $P < 0.05$) between comparisons.

**b** Summary of differentially metabolites (determined by $P < 0.05$) between comparisons.
